# Adenosine Deaminase Acting on RNA 1 Associates with Orf Virus OV20.0 and Enhances Viral Replication

**DOI:** 10.1101/459149

**Authors:** Guan-Ru Liao, Yeu-Yang Tseng, Jing-Yu Tseng, Fong-Yuan Lin, Yumiko Yamada, Hao-Ping Liu, Chih-Ying Kuan, Wei-Li Hsu

**Affiliations:** Graduate Institute of Microbiology and Public Health, National Chung Hsing University, 145 Xingda Road, Taichung, 402, Taiwan; John Curtin School of Medical Research, Australian National University, Canberra, Australian Capital Territory, Australia; Department of Beauty Science, MeiHo University, Neipu, Pingtung County, Taiwan; Department of Veterinary Medicine, National Chung Hsing University, 145 Xingda Road, Taichung, 402, Taiwan

**Author notes:** corresponding author. E-mail address (W-L Hsu).

## Abstract

Orf virus (ORFV) infects sheep and goats and is also an important zoonotic pathogen. The viral protein OV20.0 has been shown to suppress innate immunity by targeting the double-stranded RNA (dsRNA)-activated protein kinase (PKR) by multiple mechanisms. These mechanisms include a direct interaction with PKR and binding with two PKR activators, dsRNA and the cellular PKR activator (PACT), which ultimately leads to the inhibition of PKR activation. In the present study, we identified a novel association between OV20.0 and adenosine deaminase acting on RNA 1 (ADAR1). OV20.0 bound directly to the dsRNA binding domains (RBDs) of ADAR1 in the absence of dsRNA. Additionally, OV20.0 preferentially interacted with RBD1 of ADAR1, which was essential for its dsRNA binding ability and for the homodimerization that is critical for intact adenosine (A)-to-inosine (I) editing activity. Finally, the association with OV20.0 suppressed the A-to-I editing ability of ADAR1, while ADAR1 played a proviral role during ORFV infection by inhibiting PKR phosphorylation. These observations revealed a new strategy used by OV20.0 to evade antiviral responses via PKR.

**Importance:** Viruses evolve specific strategies to counteract host innate immunity. ORFV, an important zoonotic pathogen, encodes OV20.0 to suppress PKR activation via multiple mechanisms, including interactions with PKR and two PKR activators. In this study, we demonstrated that OV20.0 interacts with ADAR1, a cellular enzyme responsible for converting adenosine (A) to inosine (I) in RNA. The RNA binding domains, but not the catalytic domain, of ADAR1 are required for this interaction. The OV20.0-ADAR1 association affects the functions of both proteins; OV20.0 suppressed the A-to-I editing of ADAR1, while ADAR1 elevated OV20.0 expression. The proviral role of ADAR1 is likely due to the inhibition of PKR phosphorylation. As RNA editing by ADAR1 contributes to the stability of the genetic code and the structure of RNA, these observations suggest that in addition to serving as a PKR inhibitor, OV20.0 might modulate ADAR1-dependent gene expression to combat antiviral responses or achieve efficient viral infection.

## Introduction

Orf virus (ORFV), which belongs to the *Parapoxvirus* genus of the *Poxviridae* family, causes contagious ecthyma, mainly in sheep, goats, and other ruminants (1). In addition, humans are susceptible to ORFV and often infected by direct contact with infected animals. Previous studies suggest that ORFV encodes several gene products that can manipulate cellular antiviral responses(2). The OV20.0 protein, an ortholog of E3 from vaccinia virus (VACV), is an ORFV virulence factor, by which ORFV subverts host innate immunity and thus establishes efficient replication (3, 4). OV20.0 contains two binding domains, including a Z-DNA binding domain at the N terminus and a double-stranded RNA binding domain (RBD) at the C terminus (5). Furthermore, the *OV20.0L* gene encodes two isoforms, a full-length OV20.0 and an N-terminally truncated short form (sh20). Although the cellular distributions of these proteins differ, both OV20.0 isoforms have similar functions, such as binding dsRNA, interacting with double-stranded RNA (dsRNA)-activated protein kinase (PKR) and inhibiting PKR phosphorylation (5).

Multiple strategies that are used by OV20.0 to suppress PKR activation were revealed recently (5, 6). OV20.0 is able to associate with PKR and compete with two cellular activators of PKR, i.e., dsRNA and PACT. OV20.0 is a dsRNA binding protein (DRBP) that sequesters dsRNA and therefore prevents PKR activation (5). In addition, the interaction between OV20.0 and the RBDs of PKR arrests PKR phosphorylation by reducing the ability of dsRNA to bind to this region. This physical association also occupies the binding site for PACT in the RBD2 and kinase domains of PKR. Another possible mechanism is that OV20.0 forms heterodimers with PACT via their RBDs, which may further reduce PACT-dependent PKR activation (6).

Adenosine deaminase acting on RNA (ADAR), which is also a DRBP, converts adenosines (A) into inosines (I) in cellular and viral dsRNA through its deaminase activity. This reaction is referred to as A-to-I RNA editing (7, 8). Inosine is recognized as guanine in RNA duplexes, and A-to-I RNA editing may thus change the sequence and structure of RNA, leading to genetic recoding and the destabilization of RNA. In mammalian cells, the ADAR family includes ADAR1, ADAR2 and ADAR3; among those proteins, ADAR3 is catalytically inactive even though it contains a putative catalytic domain (9). Due to alternative promoter usage, the ADAR1 gene is transcribed into two isoforms (10) known as*p150* and *p110*; *p150* harbors a consensus sequence for the IFN-stimulated response element (ISRE) that places *p150* expression under the control of IFN (11), while *p110* is constitutively expressed. The p110 protein is an N-terminally truncated version of p150, and the two ADAR1 isoforms contain either one (i.e., p110) or two (p150) Z-DNA binding domains at the N-terminus, adjacent to three copies of the RBD (R1, R2 and R3) (12). In addition, both forms of ADAR1 have a deaminase catalytic domain at their C-terminus. Moreover, the cellular distribution of the two ADAR1 isoforms is distinct; p150 is present throughout the cell, while p110 is largely localized in the nucleus (13, 14) due to the lack of a nuclear export signal (NES).

Although ADAR1 *p150* is an IFN-inducible gene, increasing evidence indicates a proviral role of ADAR1. This protein enhances viral replication, particularly for RNA viruses, by at least two mechanisms: RNA editing and PKR inhibition (15, 16). For example, the overexpression of ADAR1 increased the yield of human immunodeficiency virus type 1 (HIV-1) in HEK 293T and COS-7 cells (17, 18); consistent with this finding, knock-down endogenous levels of ADAR1 reduced HIV protein accumulation in HEK 293T cells (18). Furthermore, ADAR1 with a catalytic domain deletion failed to enhance viral infectivity and virion release, which indicates that the proviral activity of ADAR1 is regulated by its RNA-editing ability in the context of HIV-1 infection. Independent of RNA editing, ADAR1 also elevates HIV-1 infectivity by interacting with PKR via its RBD, which leads to the inhibition of PKR phosphorylation (19). Similar to observations in HIV-1, measles virus (MV) grew poorly in cells with deficient ADAR1 expression, and that change was accompanied by the suppression of the dsRNA-mediated antiviral response, particularly PKR and IFN regulatory transcription factor-3 (IRF-3) (20). The RNA-editing activity of ADAR1 plays a role not only for RNA viruses but also for DNA viruses that replicate in the nucleus (21, 22), for example, the *Herpesviridae* family members Kaposi sarcoma-associated virus (KSHV) and Epstein-Barr virus (EBV). A-to-I editing activity eliminated the tumorigenic activity of KSHV, and affected latent infection of EBV by modifying the sequences of the viral kaposin transcript and miR-BART6, respectively (21, 22). For poxvirus, VACV E3 is a potent inhibitor of ADAR1 deaminase activity (23). In addition to two highly conserved residues in the Z-DNA binding domain, the RBD at the C-terminus of VACV E3 is essential for ADAR1 antagonism. Nevertheless, the role of ADAR1 in VACV has not been addressed.

Previous reports revealed that OV20.0 acts as a PKR antagonist by targeting its activators (5, 6). In this study, we discovered a novel interaction of OV20.0 with ADAR1 and investigated its consequences for ORFV replication. Initially, the association of OV20.0 and ADAR1 was investigated. Subsequently, a series of deletion constructs of OV20.0 and ADAR1 were made to map the domains involved in the interaction between these two proteins. Moreover, whether OV20.0 interferes with the RNA-editing activity of the ADAR1 enzyme, and the effects of ADAR1 on PKR activation were investigated. Since OV20.0 can associate with both PKR and ADAR1, the hierarchy of these associations was also examined. Finally, the impact of ADAR1 on ORFV infection was explored for the first time.

## Materials and methods

### Cell culture

Human embryonic kidney cell line 293T (HEK 293T), human lung carcinoma A549 cells, and primary goat fibroblasts were propagated in Dulbecco’s modified Eagle’s medium (DMEM; Gibco BRL, Life Technologies Corporation Carlsbad, CA, USA) with 10% fetal bovine serum (FBS; Hyclone, Logan, UT, USA) and 1% penicillin-streptomycin (Gibco BRL). All cells were incubated at 37 °C with 5% CO_2_.

### Virus and infection

The recombinant ORFV expressing enhanced green fluorescent protein (eGFP) was described previously (5). For infection, cells at 80% confluency were washed with phosphate-buffered saline (PBS) and then infected at the indicated multiplicity of infection (MOI) in DMEM without FBS. After 1 hr of incubation, the infection medium was removed and replaced with DMEM containing 2.5% FBS.

### Generation of ADAR1 knock out HEK 293T cell line

To study the role of ADAR1 play in the course of infection, CRISPR-Cas9 system was applied to inactive ADAR1 expression in HEK 293T cell line. For the ease of monitoring Cas9 expression, mCherry was added into the original plasmid, lentiCRISPR v2 (Addgene plasmid # 52961), followed by P2A sequence (derived from porcine teschovirus-1). The mCherry fragment was amplified by a set of primers (mCherry-infusion-F and mCherry-infusion-R, sequences were listed in Table 1), and the amplicon was inserted into the Tth111I site of lentiCRISPR v2 by In-Fusion cloning, resulting in the plasmid, letniCRISPR v2-mCherry. In brief, the annealed oligonucleotide containing target sequence of ADAR1, i.e. sense: 5’-GGACAGGA GACGGAATTCGC-3’ was introduced into the sites of BsmbI of lentiCRISPR v2-mCherry. HEK 293T cells were transfected with 1 μg plasmid using Lipofectamine 3000. At 48 hours post transfection, the cells were split followed by limiting dilution, which allows formation of single cell colonies. Single colony was selected for validation of ADAR1 expression by western blot analysis.

**Table 1.**
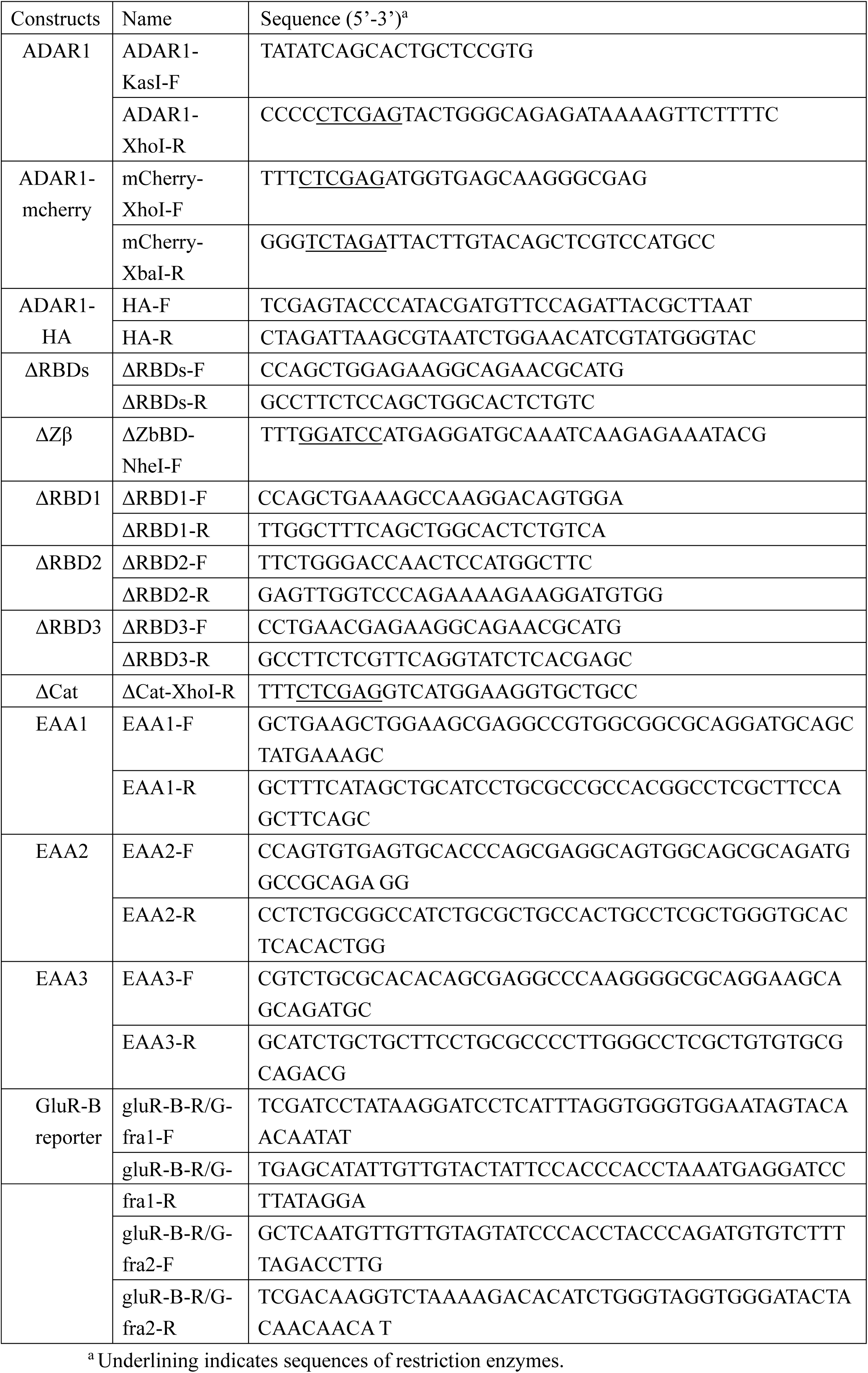
Primers used in this study.

### Construction of plasmids

The original plasmid encoding the constitutive form (p110) of ADAR1 was kindly provided by Professor Charles Samuel (Department of Molecular, Cellular, and Developmental Biology, University of California, Santa Barbara). To facilitate the pull-down experiments, a hemagglutinin (HA) tag was fused to the 3’ end of ADAR1. Briefly, to remove stop codon of ADAR1 from the original ADAR1 plasmid, the region at the 3’ end of ADAR1 in the plasmid was initially replaced with a PCR amplicon amplified by a set of primers (ADARl-KasI-F and ADARl-Xhol-R) using the original ADAR1 plasmid as a template. Subsequently, the annealed oligonucleotides of HA tag (namely, HA-F and HA-R) with compatible ends of restriction enzymes *Xho* I and *Xba* I or the entire coding sequence of mCherry amplified from mCherry N1 (Clontech, Mountain View, CA, USA) by PCR using the primers mCherry-XhoI-F and mCherry-XbaI-R was then inserted into the 3’ end of ADAR1, resulting in constructs expressing ADAR1 fused with HA tag or mCherry, respectively. The sequences of the primers and oligonucleotides used in this study were summarized in Table 1.

A set of ADAR1 deletion constructs was made by either conventional molecular cloning or In-Fusion^®^ HD cloning (Takara Bio USA, Inc. Mountain View, CA, USA). Conventional cloning was used to construct plasmids containing a deletion of the Zβ DNA binding domain (ΔZβ) and a deletion of the deaminase catalytic domain (ΔCat) of ADAR1. Fragments were amplified by designed primer sets (ΔZβ-NheI-F and HA-R; and (ΔCat)-NheI-F and (ΔCat)-XhoI-R) and then ligated into the ADAR1 vector in frame with the HA tag. Plasmids with a deletion of all double-stranded RNA binding domains (ΔRBDs) and deletions of individual RBDs (ΔR1, ΔR2 or ΔR3) of ADAR1 were separately cloned by In-Fusion^®^ HD cloning (Takara Bio USA, Inc. Mountain View, CA, USA). Fragments with the designated deletions were amplified by specific primers (ΔRBDs-F and ΔRBDs-R; ΔRBD1-F and ΔRBD1-R; ΔRBD2-F and ΔRBD2-R; and ΔRBD3-F and ΔRBD3-R) using ADAR1-p110-HA as a template. Subsequently, the coding sequences of full-length ADAR1 in the ADAR1-p110-HA plasmid were then replaced by the desired fragments, following the manufacturer’s instructions.

Moreover, the KKxxK motif in each of the three RBDs of ADAR1 was altered to the sequence EAxxA by site-directed mutagenesis using the following primer sets: EAA1-F and EAA1-R; EAA2-F and EAA2-R; and EAA3-F and EAA3-R. Constructs expressing various OV20.0 proteins fused with the FLAG tag were described previously (5).

To investigate the A-to-I editing activity of ADAR1, an mCherry reporter construct was made. Briefly, mCherry N1 (Clontech) was digested with *BamH* I and *Bgl* II, followed by self-ligation to remove one additional *Sal* I sequence in the multiple cloning site. The entire sequence of the glutamate B receptor (GluR-B) containing an in-frame amber stop codon (UAG) was assembled from two fragments, which were generated by two sets of oligonucleotides: gluR-B-R/G-fra1-F and gluR-B-R/G-fra1-R; and gluR-B-R/G-fra2-F and gluR-B-R/G-fra2-R. The two annealed GluR-B fragments with comparable sticky ends were then ligated and inserted into the *Sal* I-linearized mCherry plasmid described above.

All fragments were amplified by polymerase chain reaction (PCR) using Phusion^®^ High-Fidelity DNA Polymerase (New England Biolabs, Beverly, MA, USA). The PCR conditions were set as follows: initial denaturation at 98°C for 2 min and 35 cycles consisting of denaturation at 98°C for 10 sec, annealing at 55°C for 20 sec and extension at 72°C for 15 sec per 1 kb of the desired amplicon, followed by a final extension at 72°C for 7 min. The vectors and inserts were digested with the corresponding restriction enzymes (New England Biolabs).

### Transfection and immunoprecipitation

Briefly, HEK 293T cells were seeded 1 day before transfection. In general, for a 6-well plate scale, 5 μg of ADAR1 plasmid and 2.5 μg of OV20.0 plasmids were transfected by Lipofectamine 3000 (Invitrogen, Carlsbad, CA, USA), according to the manufacturer’s instructions. For induction of PKR phosphorylation, cells were transfected with 150 ng ploy I:C (Sigma-Aldrich, St. Louis, MO, USA). After 4 hr transfection, cells were harvested in the SDS sample dye, resolved by sodium dodecyl sulfate polyacrylamide gel electrophoresis (SDS-PAGE), and detected by Western blotting.

For immunoprecipitation (IP), 24 hr after transfection, the cells were harvested in lysis buffer (50 mM Tris [pH 7.4], 150 mM NaCl, 1 mM EDTA and 1% Triton X-100) supplemented with proteinase inhibitor cocktail (Roche Diagnostics GmbH, Mannheim, Germany) and incubated on ice for 30 min, followed by centrifugation at 13,000 rpm for 15 min. One-tenth of the supernatant was saved as the input control. The remaining cell extract was incubated with an anti-FLAG M2 affinity gel (Sigma-Aldrich, St. Louis, MO, USA) at 4°C overnight. The next day, the gel was harvested by centrifugation at 13,000 rpm, followed by ten washes with wash buffer (50 mM Tris [pH 7.4] and 150 mM NaCl). Target proteins were then eluted in SDS sample dye and separated by SDS-PAGE, followed by western blot analysis.

### dsRNA (poly I:C) pull-down assay

Agarose conjugated to poly-cytosine (Sigma-Aldrich) was mixed with 2 volumes of poly-inosine (2 mg/ml) in buffer containing 50 mM Tris (pH 7.0) and 150 mM NaCl. To form dsRNA, the mixture was gently mixed on a rotator at 4°C overnight. The agarose beads with dsRNA were then harvested by centrifugation at 1,000 rpm, washed 5 times with wash buffer (50 mM Tris [pH 7.0] and 150 mM NaCl), and resuspended in binding buffer (50 mM Tris [pH 7.5], 150 mM NaCl, 1 mM EDTA, and 1% NP-40).

The cell lysate expressing the desired proteins was mixed with annealed poly I:C beads on a rotator at 4°C overnight. The next day, the poly I:C beads were collected by centrifugation at 1,000 rpm and then washed ten times with wash buffer. Proteins associated with the dsRNA were eluted in SDS sample dye.

### Western blot analysis

The samples were separated by SDS-PAGE and then transferred to nitrocellulose paper (NC paper). After blocking in 5% skim milk, the primary antibodies were incubated with the NC paper at 4°C overnight. The next day, the NC papers were washed with PBS plus 0.05% Tween 20 five times, followed by incubation with the corresponding secondary antibody conjugated to horseradish peroxidase (HRP). Protein detection was performed by an ImageQuant LAS 4000 (GE Healthcare, Uppsala, Sweden). The corresponding dilutions of each antibody were as follows: anti-ADAR1 (1:1,000; Santa Cruz Biotechnology), anti-PKR (1:2,000, Abcam), anti-PKR-p (T446; 1:2,000; Abcam), anti-β-actin (1:1,000; Signalway Antibody), anti-GAPDH (1:1,500; Novus Biologicals), anti-FLAG (1:2,500; Signalway Antibody), anti-HA (1:2,500; Yao-Hong Biotechnology) and anti-0V20.0 (1:2,000) (5).

### Immunofluorescence assay (IFA)

The transfected cells were fixed with 1.875% formaldehyde, permeabilized with 0.5% NP-40, and then incubated with an ADAR1 antibody (sc-73408, 1:100 dilution; Santa Cruz Biotechnology) for 1 hr at room temperature. The cells were further incubated with secondary antibodies conjugated to Alexa Fluor 488 (1:5,000 dilution, Invitrogen) for additional 1 hr at room temperature. Wash procedures were applied between each step, with six washes with 1% FBS in PBS, followed by DAPI (4’, 6-diamidino-2-phenylindole, at a final concentration of 1 μg/ml, Invitrogen) staining. The cells were examined by an inverted fluorescence confocal microscope (FV1000, Olympus, Tokyo, Japan). Images were acquired by Olympus FV10-ASW 1.3 viewer software.

## Results

### OV20.0 interacts with ADAR1 via its RBD

0V20.0 regulates antiviral responses by interacting with several cellular DRBPs, including PKR and PACT (5, 6). Since DRBPs tend to form homodimers or heterodimers with other DRBPs (24), we suspected that 0V20.0 may associate with other DRBPs and influence the fate of virus infection. ADAR1, a cellular DRBP, interacts with PKR and leads to different consequences during viral replication (15). Unlike p110, p150 ADAR1 was not obviously induced in ORFV infection (data not shown), only p110 was represented in this study. To determine the binding domain involved in the OV20.0-ADAR1 interaction, a set of OV20.0 plasmids was constructed to express the wild-type form (WT, encoding two isoforms), the full-length isoform only (K20), the N-terminally truncated short form (sh20), or the C-terminally truncated form (ΔC) of OV20.0 (Fig. 1A). OV20.0 and all of its variants were detected by a FLAG antibody (Fig. 1B; input). Moreover, both isoforms (K20 and the N-terminal truncation sh20) could pull down endogenous ADAR1 (Fig. 1B; left panel, IP), while the ΔC deletion abrogated the ADAR1-binding ability of the protein. These results indicated that the C-terminal RBD of OV20.0 was required for the interaction between OV20.0 and ADAR1, while the Z-DNA binding domain at the N-terminus was dispensable.

**Fig 1.**
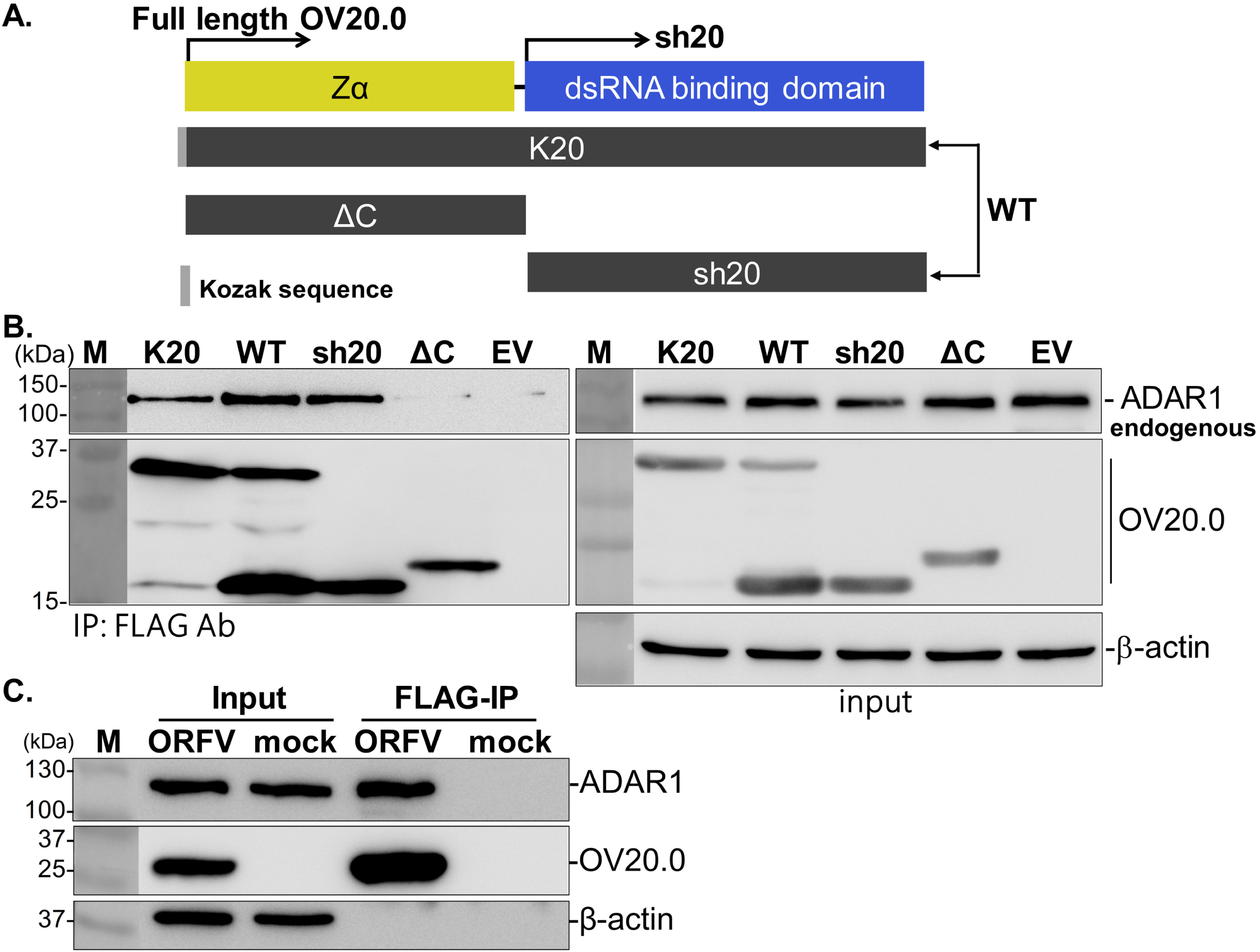
Interaction of ORFV OV20.0 with ADAR1. (A) A set of constructs expressing the wild-type form and deletions of ORFV OV20.0 were used in this study. With the Kozak sequence modification, full-length OV20.0 is preferentially expressed. The interaction of OV20.0 with endogenous ADAR1 in a transient expression system (B) and in ORFV-infected cells (C) was determined by an immunoprecipitation assay (IP). (B) Plasmids expressing FLAG-tagged wild-type OV20.0, Kozak OV20.0 (K20), short form OV20.0 (sh20), C-terminus deletion OV20.0 (ΔC), and FLAG empty vector (EV, as a negative control for the IP procedure) were transfected individually into HEK 293T cells, followed by IP with an FLAG-antibody (FLAG-IP). (C) HEK 293T cells were infected with a recombinant eGFP reporter ORFV expressing FLAG-tagged OV20.0 at an MOI of 1 for 24 hr, followed by FLAG-IP.

Next, a recombinant ORFV expressing FLAG-tagged OV20.0 (5) was used to verify the association between OV20.0 and ADAR1 in the context of ORFV infection. HEK 293T cells were infected with ORFV-FLAG at an MOI of 1, and the cell lysate was harvested at 24 hr post infection (hpi), followed by FLAG-IP. Consistent with results of transient expression (Fig. 1B), endogenous ADAR1 was associated with FLAG-tagged OV20.0 produced during ORFV infection (Fig. 1C).

### Cellular distribution of OV20.0 and ADAR1

Cellular distribution of OV20.0 isoforms has been described (5); K20 is a nuclear protein, and sh20 expresses in both nuclear and cytoplasm. As OV20.0 associates with ADAR1, whether that protein-protein interaction can lead to an alteration of protein cellular distributions is of interest. To monitor subcellular localization, ADAR1 and OV20.0 were expressed with an mCherry or eGFP tag at their C-terminus, respectively. As observed for endogenous ADAR1, ADAR1-mCherry was expressed as a speckled pattern in the nucleus (left and right panels in Fig. 2A, respectively). Further, the integrity of the ADAR1-mCherry fusion protein and the expected size of approximately 150 kDa were confirmed by western blot analysis (Fig. 2A, lane 2 indicated with an arrow).

**Fig 2.**
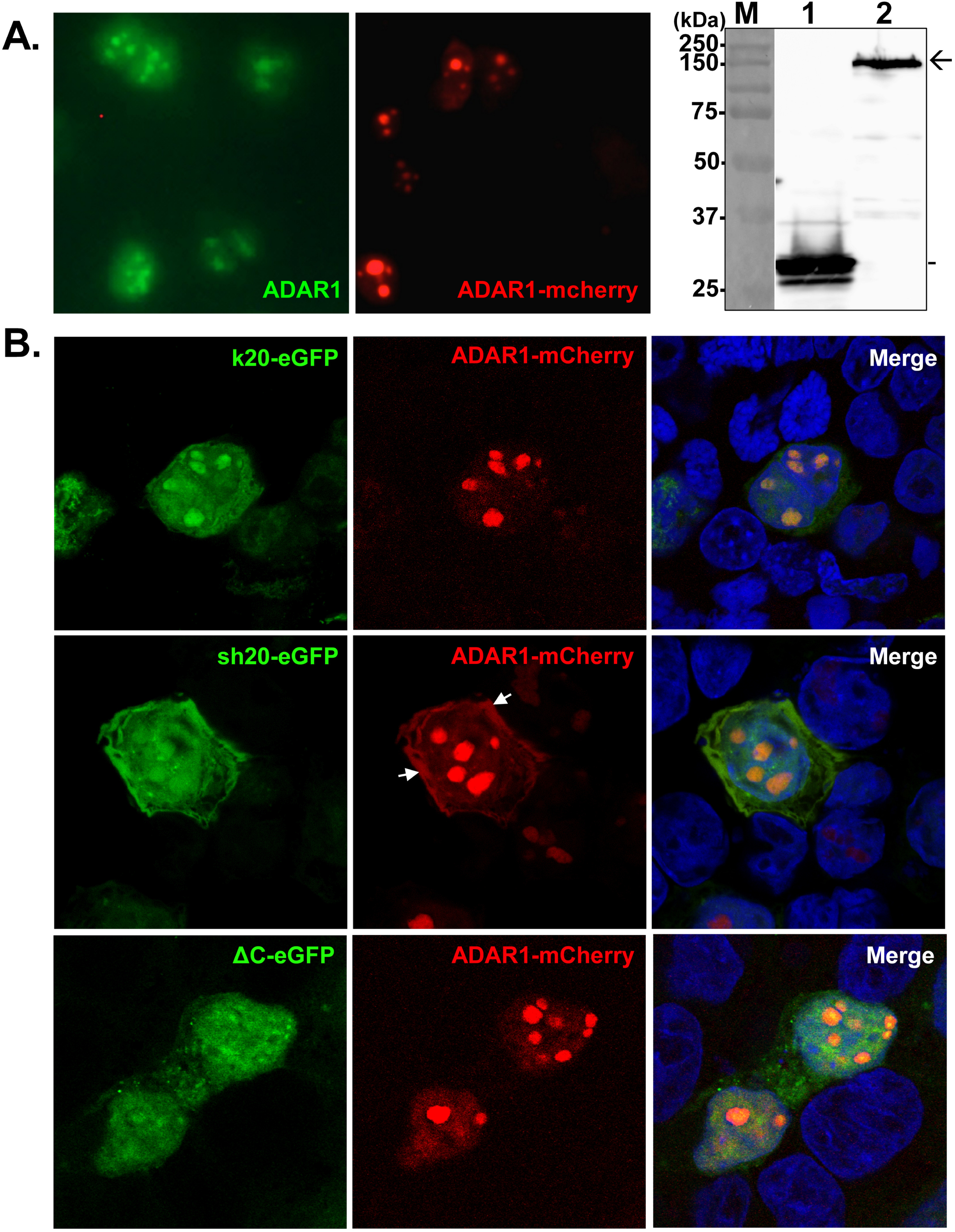
Cellular distribution of ADAR1 in the presence of OV20.0. The expression of endogenous ADAR1 and ADAR1 fused with mCherry was determined by an immunofluorescence assay using an ADAR1 antibody or autofluorescence, respectively. The molecular weight and integrity of mCherry (lane 1) and ADAR1-mCherry (lane 2) were determined by western blot analysis using an mCherry antibody. (B) Colocalization of ORFV OV20.0 isoforms with ADAR1. HEK 293T cells were cotransfected with one of the eGFP-tagged OV20.0 variants (K20, sh20, or ΔC) and the ADARl-mCherry constructs. The cellular distribution of ADAR1 and OV20.0 was examined by fluorescence microscopy. White arrows indicate cytoplasmic distribution of ADAR1.

When K20-eGFP was expressed with ADAR1-mCherry, the proteins colocalized in the nucleus (Fig. 2B). Approximately, ADAR1 colocalization was observed in 50% (53 out of 106), or 64% (106 out of 166) of cells overexpressing K20-eGFP or sh20-eGFP, respectively. Interestingly, in the presence of sh20-eGFP, a portion of the ADAR1-mCherry signal shifted from the nucleus to the cytoplasm and colocalized with sh20-eGFP in the cytoplasm (Fig. 2B, indicated with white arrows). In contrast, ΔC-eGFP was expressed throughout the cells; however, it was not particularly colocalized with ADAR1 in the nucleus. Collectively, both K20 and sh20 colocalized with ADAR1 in a transient expression system in HEK 293T cells.

### RBD1 of ADAR1 is essential for the interaction with OV20.0

To identify the region of ADAR1 that contributes to the OV20.0 interaction, a set of ADAR1 constructs with deletion of the main functional domains (Fig. 3A), including the Zβ DNA binding domain (ΔZβ), all RBDs (ΔRBDs), and the deaminase catalytic domain (ΔCat) were generated. The results of FLAG-IP showed that in the absence of the Zβ or catalytic domains, ADAR1 still associated with OV20.0. However, when ADAR1 lacked all RBDs, the interaction between OV20.0 and ADAR1 was greatly reduced (Fig. 3B, left panel).

**Fig 3.**
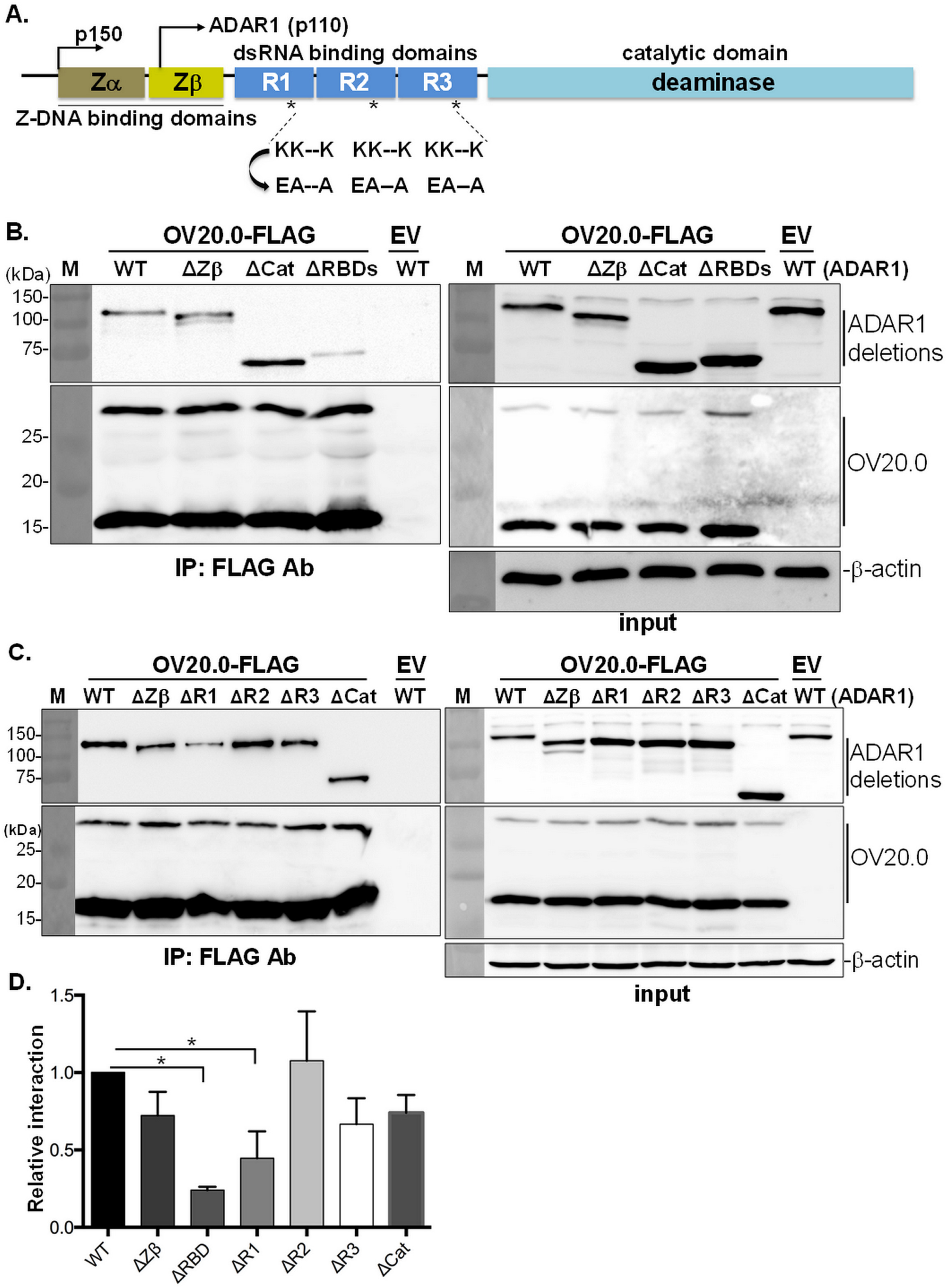
Regions of ADAR1 involved in the OV20.0 interaction. (A) The structure of ADAR1 is schematically illustrated. ADAR encodes two isoforms, namely, p150 and p110. ADAR1 contains two Z DNA binding domains (Zα and Zβ), three RBDs (indicated as R1, R2, and R3), and the deaminase catalytic domain. Mutation of the canonical KKxxA motifs in the three RBDs to EAxxA abolished the dsRNA binding ability of ADAR1. (B) The RBDs of ADAR1 are critical for the OV20.0 interaction. Plasmids expressing HA-tagged WT ADAR1 or individual deletions of the Zβ binding domain (Δ Zβ), all RBDs (Δ RBDs), or the catalytic domain (ΔCat) were cotransfected with FLAG-tagged OV20.0 (or FLAG empty vector, as a negative control for FLAG-IP) into HEK 293T cells, followed by FLAG-IP. (C) OV20.0 preferentially binds R1 of ADAR1 (C-D). To further explore the contribution of individual RBDs, constructs expressing deletions of one RBD (ΔR1, ΔR2, and ΔR3) were transfected along with FLAG-tagged OV20.0 into HEK 293T cells, followed by FLAG-IP. The IP experiment was performed with three independent repeats, and the relative interaction of ADAR1 with OV20.0 was then estimated. (D) The intensity of ADAR1 variant pull-down by FLAG-IP was initially normalized to β-actin. The interaction level of WT ADAR1 with OV2.0 was arbitrarily set as 1. All the experiment was conducted in triplicate. Note: * indicate p value < 0.05.

Next, we further examined which RBD domain in ADAR1 is important for the interaction. OV20.0 with a FLAG tag and ADAR1 with deletions of individual RBDs (ΔR1, ΔR2, or ΔR3) were transiently expressed in HEK 293T cells, followed by FLAG-IP. The data demonstrated that the deletion of R2 of the three RBDs did not have effect on the ADAR1-OV20.0 interaction (Fig. 3C). However, the absence of the first RBD of ADAR1 significantly decreased the binding between OV20.0 and ADAR1 (Fig. 3D). In conclusion, the RBDs of ADAR1, especially the first RBD, were crucial for the OV20.0-ADAR1 interaction.

### dsRNA binding ability of ADAR1 is not necessary for the interaction between OV20.0 and ADAR1

According to a previous report, OV20.0 interacts with and disrupts the function of other DRBPs, such as PKR, in the absence of dsRNA (6). We next investigated whether the dsRNA binding ability of ADAR1 is involved in the interaction with OV20.0. Therefore, an ADAR1 mutant (EAA) that is defective in dsRNA binding ability (25, 26) was used to address this issue. A triple-nucleotide point mutation (EAxxA) was introduced in each RBD of ADAR1 to replace the original dsRNA binding motif (KKxxK). First, as with previous findings (25, 26), the EAxxA mutations in the three RBDs indeed impeded the dsRNA binding ability of ADAR1 in a poly I:C pull-down assay (Fig. 4A). Then, FLAG-IP was performed to investigate the interaction between OV20.0 and the EAA mutant ADAR1; the data showed that OV20.0 associated with WT ADAR1 and the EAA mutant to similar extents (Fig. 4B). In conclusion, we found that the dsRNA binding ability of ADAR1 is not essential for the interaction of ADAR1 with OV20.0.

**Fig 4.**
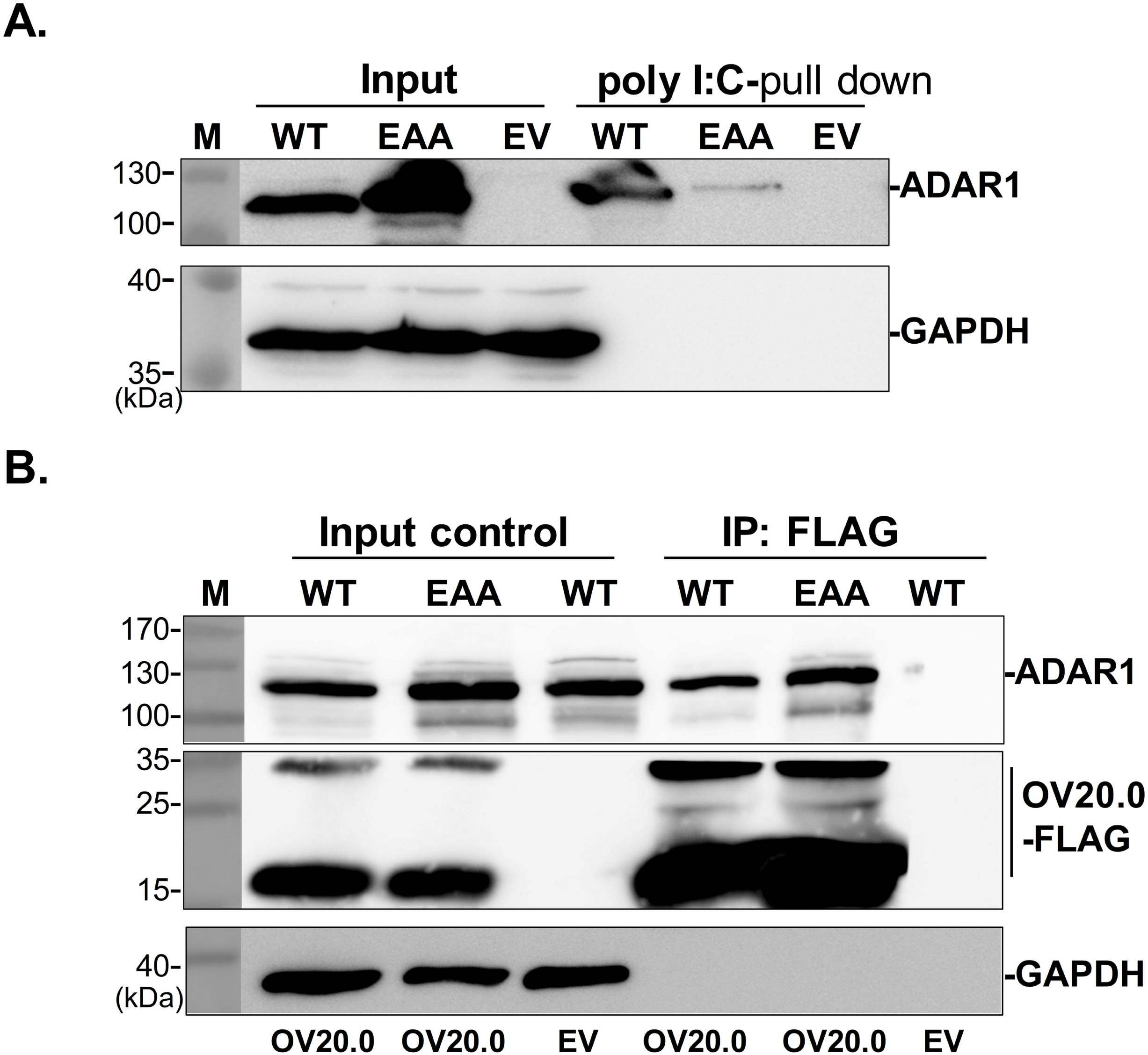
dsRNA binding ability is not required for the interaction of OV20.0 and ADAR1. (A) Plasmids expressing HA-tagged ADAR1 (WT), dsRNA binding-deficient ADAR1 (with EAA mutations in the three RBDs; labeled EAA), and HA empty vector were transfected individually into HEK 293T cells, followed by a poly I:C (as a synthetic dsRNA) pull-down assay. (B) Plasmids expressing HA-tagged WT ADAR1, an EAA mutant, or the FLAG empty vector were cotransfected with FLAG-tagged OV20.0 into HEK 293T cells, followed by FLAG-IP.

### OV20.0 disrupts the A-to-I editing activity of ADAR1

The A-to-I editing activity of ADAR1 has been shown to enhance or prevent viral replication in a cell- and virus-dependent manner; however, it has only been characterized for a few viruses (27). At present, the impact of the association of OV20.0 on ADAR1 remains unclear. To address this issue, an mCherry reporter assay was established to measure the A-to-I editing activity of ADAR1. The sequence of the glutamate B receptor (GluR-B) containing an in-frame amber stop codon (UAG) (28) was inserted into the mCherry coding region near its C-terminus, resulting in the translation of a C-truncated product (Fig. 5A, unedited). In the presence of ADAR1 A-to-I editing, the stop codon U<u>A</u>G could be converted to U<u>I</u>G, leading to the expression of full-length mCherry (Fig. 5A, edited). As shown in Fig 5B, in the absence of ADAR1, only truncated mCherry was expressed (Fig 5B, lane 1), while ADAR1 over-expression led to the translation of full-length mCherry in a dose-dependent manner (Fig. 5B, lane 2-4).

**Fig 5.**
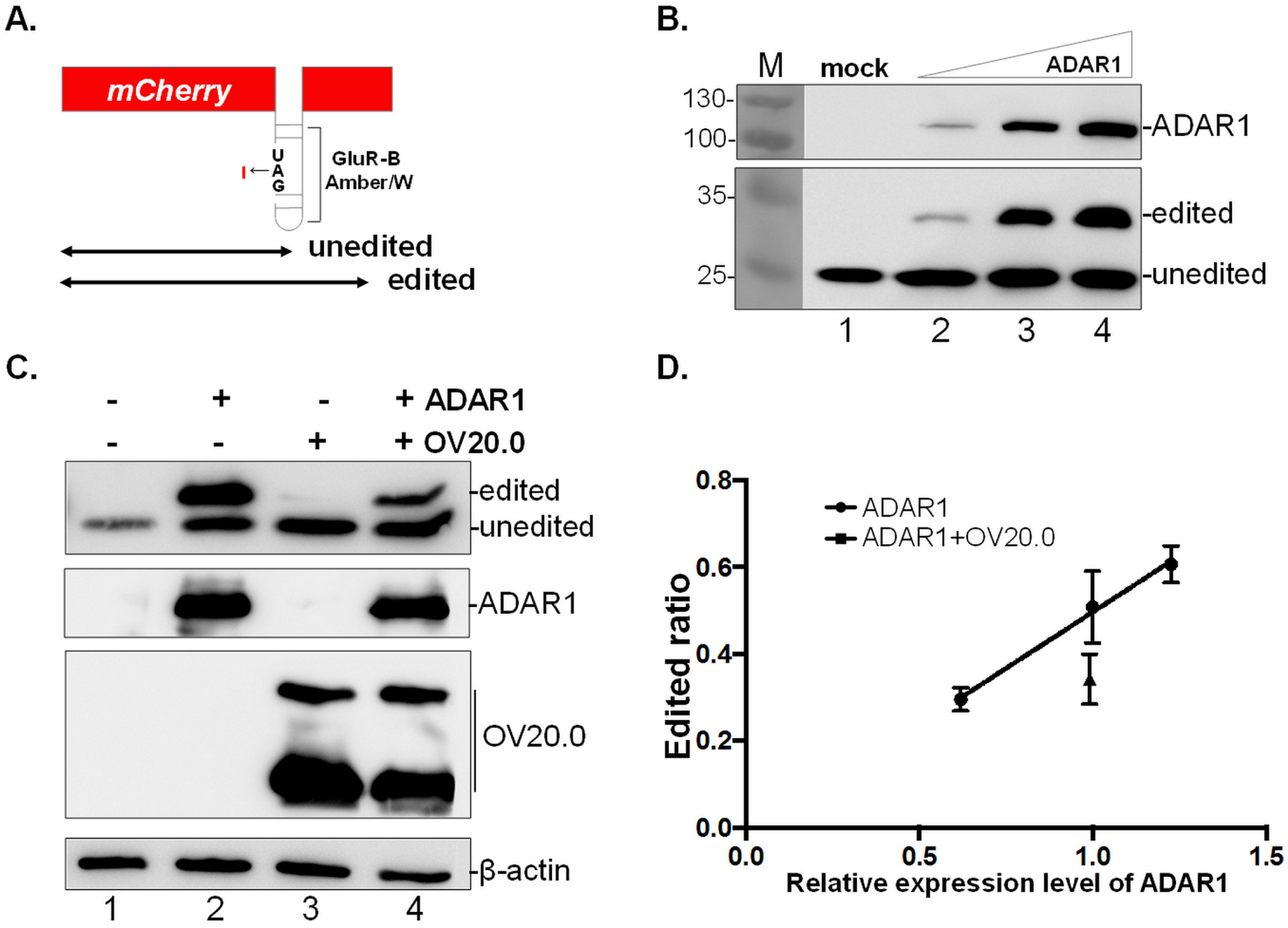
OV20.0 interferes with ADAR1-dependent A-to-I editing activity. (A) An illustration of the mCherry-based reporter system for monitoring A-to-I editing activity. The GluR-B Amber/W editing site, which contains target sequences for ADAR1-mediated RNA editing, was inserted near the C-terminus of the mCherry coding region. The built-in UAG codon insertion leads to the translation of a truncated version of mCherry, while the ADAR1 enzyme likely converts A to I to yield a full-length mCherry. (B) The ADAR1 enzyme edits the sequence of the reporter transcript in a dose-dependent manner. HEK 293T cells were transfected with the reporter plasmid, along with increasing amounts of HA-tagged ADAR1 (0, 0.2, 0.4, and 0.8 μg). At 24 hr after transfection, A-to-I editing activity was assessed by western blot analysis using antibodies against HA and mCherry. (C) The effect of OV20.0 on ADAR1-mediated A-to-I editing. HEK 293T cells were cotransfected with the mCherry reporter plasmid and HA-tagged ADAR1 alone or with FLAG-tagged OV20.0 for 24 hr. (D) A standard curve indicating the effect of A-to-I editing activity was established by measuring the ratio of edited/unedited products with increasing amounts of ADAR1. The experiment was conducted with three independent repeats.

Compared with ADAR1 alone (Fig. 5C, lane 2), the coexistence of OV20.0 significantly reduced the A-to-I editing activity of ADAR1 (lane 4 in Fig 5C, and Fig 5D). Noticeably, the expression of OV20.0 alone had no effect on A-to-I editing (Fig 5C, lane 3). These results suggested that OV20.0 interrupted the A-to-I editing activity of ADAR1.

### ADAR1 plays a proviral role in ORFV infection

To date, the role of ADAR1 in ORFV replication remains unknown. Moreover, both ADAR1 and OV20.0 interfere with PKR activation, and it is of interest to investigate the effect of ADAR1 on PKR signaling in the presence of viral OV20.0.

First, the effect of ADAR1 on PKR phosphorylation was verified. Compared with mock treatment, PKR activation was indeed stimulated by the transfection of synthetic dsRNA, poly I:C (Fig. 6A, lane EV). However, PKR activation was lower in cells overexpressing ADAR1 than in cells transfected with empty vector (EV) (Fig. 6A, right panel). Moreover, the deletion of the RBDs slightly restored PKR activation compared with WT ADAR1 (Fig. 6B). Next, since OV20.0 and ADAR1 associate with each other and both proteins are able to inhibit PKR phosphorylation, we further investigated whether the presence of OV20.0 influences the inhibitory effect of ADAR1 on PKR activation. As shown in Fig 6C, compared with the positive control (PC), OV20.0 suppressed PKR phosphorylation (PC vs. lane 1), and the coexistence of ADAR1 further inhibited PKR activation (Fig. 6C, lane 1 vs. lane 2-4).

**Fig 6.**
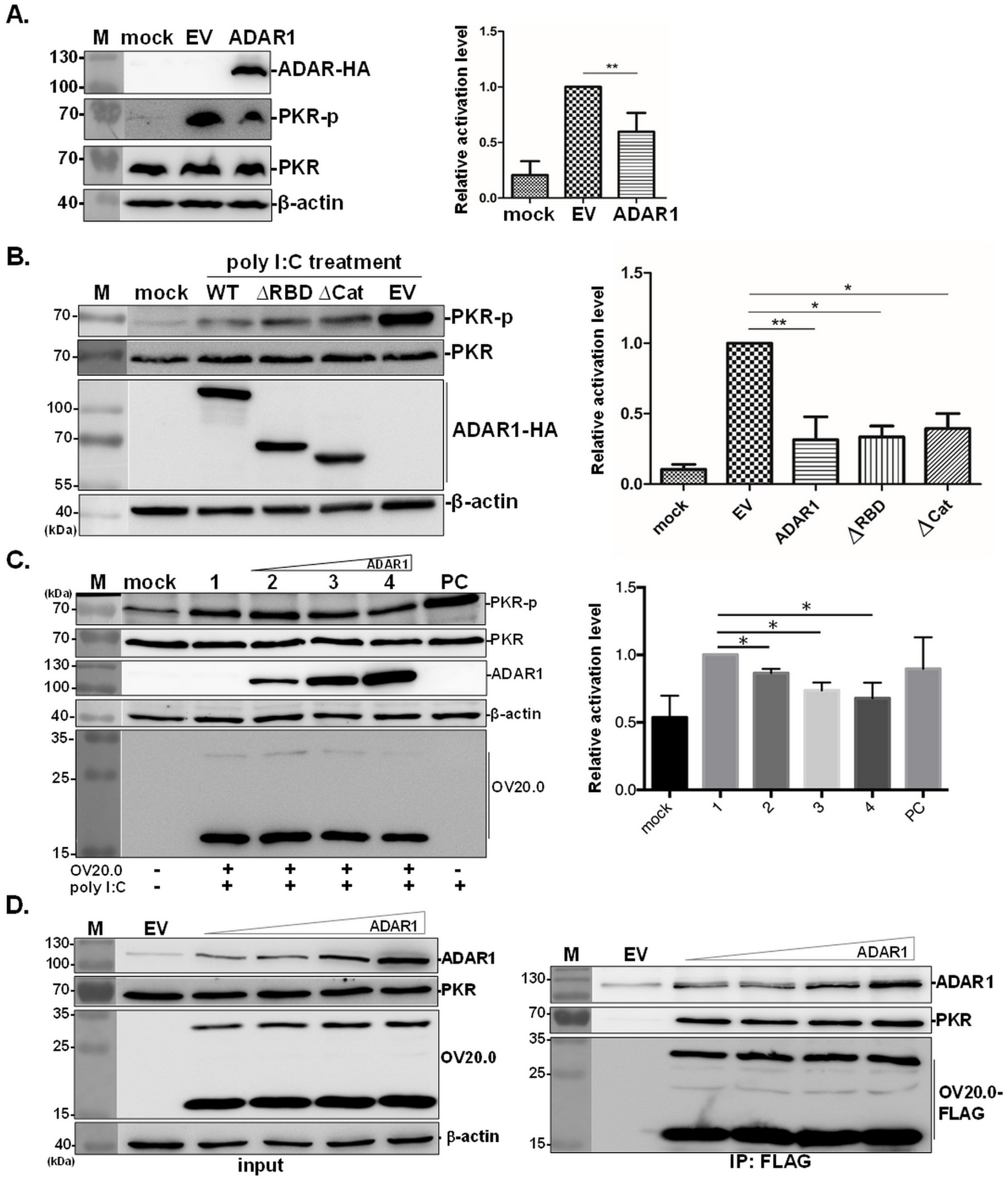
Effect of ADAR1 on PKR activation. (A) ADAR1 reduces PKR activation. HEK 293T cells were transfected with empty vector (EV) or HA-tagged ADAR1 (ADAR1), followed by activation of PKR by the transfection of 150 ng/ml poly I:C for 4 hr. The phosphorylation level of PKR was monitored by western blot analysis. The relative PKR activation level was estimated and plotted (right panel). The PKR activation level in cells transfected with EV was arbitrarily set as 1. (B) Region required for PKR suppression. HEK 293T cells were transfected with empty vector (EV) or various ADAR1 plasmids expressing the WT protein, a deletion of the three RBD (ΔRBDs) or a deletion of the catalytic domain (ΔCat) for 24 hr, and PKR phosphorylation was then induced by poly I:C. (C) ADAR1 diminishes PKR activation in the presence of OV20.0. HEK 293T cells were cotransfected with OV20.0 (0.2 μg) and increasing amounts (0.2, 0.4, and 0.8 μg) of plasmids expressing ADAR1 overnight, followed by the induction of PKR activation. Untransfected cells under poly I:C stimulation serve as a positive control (PC) for PKR activation, and mock indicates cells without any treatment. (D) ADAR1 did not compete with PKR for the interaction with 0V20.0. HEK 293T cells were cotransfected with FLAG-tagged 0V20.0 (0.2 μg) with either EV or increasing amounts of the ADAR1 construct (0.1, 0.2, 0.4, and 0.8 μg), followed by FLAG-IP and immunoblotting. All the experiment was conducted in triplicate. Note: *, ** indicate p value < 0.05, or <0.001, respectively.

Furthermore, as OV20.0 associated with both PKR and ADAR1, it is unclear whether there is a binding hierarchy of OV20.0 with these two cellular proteins. To address this question, OV20.0-FLAG was coexpressed with increasing amounts of ADAR1, followed by FLAG-IP. The results demonstrated that similar amounts of PKR interacted with OV20.0, even with gradually increasing ADAR1 levels (Fig. 6D). This finding suggested that ADAR1 did not compete with the interaction between OV20.0 and PKR.

Finally, we investigated the impact of ADAR1 on ORFV infection. ADAR1-HA or ADAR1-mCherry and the HA or mCherry vector, as a negative control, were transfected into HEK 293T cells for 24 hr, followed by infection with recombinant ORFV expressing eGFP (5). The results showed that the overexpression of WT ADAR1 led to increased virus-encoded eGFP levels (Fig. 7A). Interestingly, the elevated virus-encoded protein was significantly diminished when the cells expressed ADAR1 with a deletion of all three RBDs (Fig. 7A, right panel), suggesting that ADAR1 played a supportive role through the contribution of its RBDs during ORFV infection.

**Fig 7.**
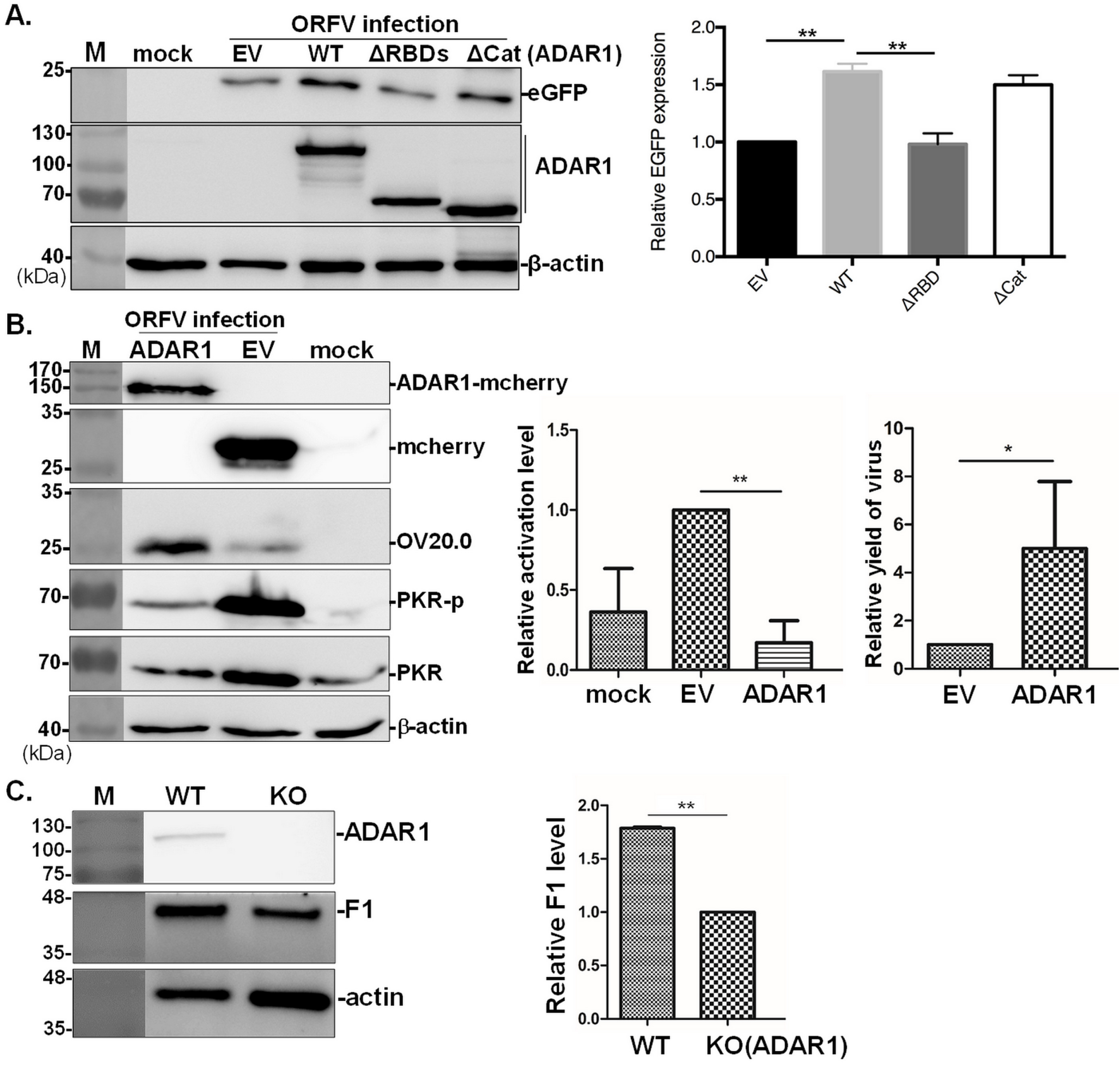
ADAR1 plays a proviral role in ORFV infection. (A) Overexpression of ADAR1 enhanced viral protein accumulation during ORFV infection. HEK 293T cells were transiently transfected with ADAR1 plasmids expressing the wild-type (WT) protein, a deletion of the three RBDs (ΔRBDs), or a deletion of the catalytic domain (ΔCat). EV indicates empty vector, which serves as a negative control. Subsequently, the transfected cells were infected with a recombinant ORFV expressing eGFP at an MOI of 1 for 48 hr, followed by immunoblotting. The experiment was conducted in triplicate, and the relative expression level of virus-encoded eGFP was estimated and plotted (right panel). The eGFP expression level in cells transfected with EV was arbitrarily set as 1. (B) The proviral role of ADAR1 is correlated with its inhibitory effect on PKR activation. HEK 293T cells transiently expressing ADAR1-mCherry (ADAR1) or mCherry (EV, as a negative control) were infected with ORFV at an MOI of 1 for 48 hr, followed by immunoblotting (left panel).The relative activation level of PKR (PKR-p) was normalized to the PKR basal level and plotted. The PKR activation level in cells transfected with EV was arbitrarily set as 1 (middle panel), and the yield of viral progenies was measured by standard plaque assay (right panel). (C) ORFV infection in ADAR1 knock out cells. 293T cells with endogenous (WT), or without (KO) ADAR1 expression were infected with ORFV Expression of viral structure protein F1 was determined at 48 hpi by western blot analysis (left panel). The relative expression level of F1 was plotted (right panel). All the experiment was conducted in triplicate. Note: *, ** indicate p value < 0.05, or <0.001, respectively.

Despite various effects of ADAR1 on virus replication, it has been reported ADAR1 enhances viral infection through the inhibition of PKR activity (16). In cells overexpressing ADAR1 during ORFV infection, we monitored the activation of PKR signaling and noticed that increased viral OV20.0 levels are correlated with a significantly lower PKR activation status (Fig 7B, middle middle). Overall, with ADAR1 over-expression, the viral yield of ORFV was increased by approximately 4 folds (Fig 7B, right panel). To further confirm the role of ADAR1, 293T cells with ADAR1 expression was knocked out by means of CRISPER technique. As shown in Fig 7C, in the absence of ADAR1 expression (KO), ORFV structural protein F1 level was lower than that in cells with endogenous ADAR1 (WT).

## Discussion

This study identified ADAR1 as a novel dsRNA binding protein (DRBP) that interacts with OV20.0 and enhances the replication of ORFV. According to previous reports, the RBD of OV20.0 mediates its interactions with both PKR and PACT, leading to the inhibition of PKR activation (5, 6). Similarly, OV20.0 and ADAR1 interact with each other via their RBD regions. In particular, the first RBD domain in ADAR1 (R1) is not only critical for the interaction with OV20.0 but also essential for the dimerization of ADAR1 (29). It is possible that OV20.0 association impairs the formation of the ADAR1 homodimer, which is the functional conformation (e.g., A-to-I editing activity). By using the EAA mutant ADAR1 with deficient dsRNA binding ability (25, 26), we further demonstrated that dsRNA binding ability of ADAR1 is not required for the OV20.0-ADAR1 interaction, which is consistent with the scenario reported for the PKR and PACT interaction (6). Similarly, OV20.0 directly interacts with PKR without the aid of dsRNA (6). Moreover, we found that ADAR1 is not able to compete with PKR for the interaction of OV20.0, implicating that OV20.0 favors binding to PKR over ADAR1. However, whether these interactions are bridged via either PKR or ADAR1 requires further investigations due to the multiple interactions among these DRBPs, OV20.0, PKR, and ADAR1.

OV20.0 was expressed as two isoforms, and both full-length OV20.0 and its N-terminally truncated short form (sh20) colocalized with ADAR1 (Fig. 2B). In our previous report, sh20 was mainly expressed in the cytoplasm due to the lack of a putative nuclear localization signal (NLS).Noticeably, a small portion of sh20.0 was colocalized with ADAR1 in nuclear domains, possibly nucleoli. Although ADAR1 largely remains in the nucleus, a dynamic subcellular distribution of ADAR1 has been reported; the protein shuttles between the cytoplasm, nucleus, and nucleolus (30, 31). It is worth noting that OV20.0 does not contain a nucleolus localization signal. Since ADAR1 localizes in the nucleolus (31–33), it is likely that the redistribution of both isoforms into subnuclear domains could be mediated by the association with ADAR1 (Fig 2B). Furthermore, ADAR1 is also translocated from the nucleus to the cytoplasm in a subset of cells coexpressing sh20 (Fig. 2), suggesting that intermolecular interactions may affect the localization of both ADAR1 and sh20. Barrauda and colleagues reported that the nuclear transportation of ADAR1 is aided by the nuclear import receptor transportin-1 (TRN1), which interacts with the region where the NLS overlaps with the RBD (namely, the RBD-NLS) of ADAR1 (34). Therefore, we postulate that the association of sh20 with ADAR1 might interrupt the binding between the RBD-NLS of ADAR1 and TRN1, which in turns leads to the cytoplasmic retention of ADAR1.

ADAR1 modulates host innate immunity against viral pathogens via its RNA-editing activity and that in turns inhibits PKR activation (15). Very recently, Chung and colleague further identified that not only dsRNA binding, also catalytic domain of ADAR1 is required for efficiently downregulates hyperactivation of PKR induced by endogenous dsRNA(26). In addition, the contribution of ADAR1 on RNase L-mediated apoptosis triggered by dsRNA was also reported (4). On the other hand, A-to-I editing activity potentially alters the genetic code or disrupts RNA structures, which can lead to either enhanced or reduced virus growth (35). For instance, several viral DRBPs, including nonstructural protein 1 (NS1) of influenza A virus and NS3 of Dengue virus, were found to interact with ADAR1 and improve its A-to-I editing activity, which is correlated with enhanced viral replication (36). Nonetheless, the RNA replication of human hepatitis C virus (HCV) was increased in ADAR1-deficient cells, indicating that ADAR1 plays a role in limiting viral infection by specifically eliminating HCV RNA (37). Accordingly, the role of ADAR1 in viral infection as either an antiviral or proviral factor is dependent on the involved viruses and cell types (27). Previous reports found that the formation of a homodimer of ADAR1 via its RBDs is important for its RNA-editing activity (25). Moreover, mutations of key residues in the third RBD (R3) in ADAR1 impaired both dsRNA-binding ability and catalytic activity (38). Collectively, these findings demonstrate that the functional conformation, dsRNA-binding ability, and deaminase activity of ADAR1 are key components that contribute to the A-to-I editing of ADAR1. The study presented herein demonstrated that OV20.0 suppresses ADAR1-dependent A-to-I editing activity (Fig 5). Since the RBDs of ADAR1 are associated with and therefore preoccupied by 0V20.0, the suppression of ADAR1 RNA editing activity could be due to the loss of its RNA binding ability or/and the inability to form functional homodimers. Furthermore, OV20.0, as a dsRNA binding protein, could compete for dsRNA substrate with ADAR1.

Vaccinia virus E3, an ortholog of OV20.0, suppressed the A-to-I editing of ADAR1 (23); however, at present, the potential role of ADAR1 in the replication of vaccinia virus, ORFV, or other members of the *Poxviridae* remains unclear. In addition to its dsRNA editing function, ADAR1 influences viral replication by interacting with PKR, which leads to the inhibition of its phosphorylation (16, 19). For efficient replication, many viruses encode gene products that target PKR or its downstream effector, illustrating the importance of PKR signaling for antiviral defense (39). ADAR1 inhibits PKR via several strategies, including the sequestration of dsRNA and the prevention of PKR activation through a direct interaction, and hence exerts a proviral activity (19, 20, 30). According to one previous report (19) and our data, the RBDs of ADAR1 are required for suppressing PKR phosphorylation. This observation suggests that ADAR1 competes for binding of the PKR activator, dsRNA, via its RBDs, which are also a potential domain by which ADAR1 contacts and forms an inactive dimer with PKR, leading to the inhibition of PKR activation. The effect of ADAR1 on PKR activation is well studied in the context of retrovirus infection (19, 40). It was shown that both full-length ADAR1 and a mutant ADAR1 with a catalytic domain deletion, rather than a truncation mutant in the RBDs, can restore HIV production inhibited by PKR. The results demonstrated that the attenuation of the inhibitory effect of PKR on HIV requires the RBDs of ADAR1 but not its deaminase function (19). Consistent with this finding, another study showed that the expression of wild-type and editing-deficient ADAR1 leads to comparable increases in human T-cell leukemia virus (HTLV) release and viral protein expression (40). Further, the level of PKR activation is inversely correlated with the level of ADAR1 expression. Collectively, the proviral effect of ADAR1 on HTLV is mediated by the inhibition of PKR phosphorylation and is independent of its editing activity. These data suggest that the impact of ADAR1 on viral replication could be variable owing to the versatile activity of ADAR1 (an RNA modifier, a DRBP) and the dynamic interactions among the DRBPs (PKR, ADAR1, and viral proteins).

In conclusion, several novel findings were revealed in this study. First, the association of OV20.0 and ADAR1 was identified for the first time. Moreover, this interaction affects both ADAR1 bioactivities and 0RFV infection. In particular, OV20.0 suppresses the dsRNA editing function and alters the cellular distribution of ADAR1. 0n the other hand, ADAR1 plays a proviral role in 0RFV infection, possibly via the attenuation of the antiviral effect of PKR.

## Acknowledgment

The project was supported by the Ministry of Science and Technology, Taiwan (M0ST-106-2321-B-005-039-MY3). We are grateful for the important reagents provided by Professor Charles Samuel (Department of Molecular, Cellular, and Developmental Biology, University of California, Santa Barbara, USA) and also his constructive comments during the preparation of this manuscript.

